# ABLE: an Activity-Based Level Set Segmentation Algorithm for Two-Photon Calcium Imaging Data

**DOI:** 10.1101/190348

**Authors:** Stephanie Reynolds, Therese Abrahamsson, Renaud Schuck, P. Jesper Sjöström, Simon R. Schultz, Pier Luigi Dragotti

## Abstract

We present an algorithm for detecting the location of cells from two-photon calcium imaging data. In our framework, multiple coupled active contours evolve, guided by a model-based cost function, to identify cell boundaries. An active contour seeks to partition a local region into two subregions, a cell interior and exterior, in which all pixels have maximally ‘similar’ time courses. This simple, local model allows contours to be evolved predominantly independently. When contours are sufficiently close, their evolution is coupled, in a manner that permits overlap. We illustrate the ability of the proposed method to demix overlapping cells on real data. The proposed framework is flexible, incorporating no prior information regarding a cell’s morphology or stereotypical temporal activity, which enables the detection of cells with diverse properties. We demonstrate algorithm performance on a challenging mouse *in vitro* dataset, containing synchronously spiking cells, and a manually labelled mouse *in vivo* dataset, on which ABLE achieves a 67.5% success rate.

**Significance statement:** Two-photon calcium imaging enables the study of brain activity during learning and behaviour at single-cell resolution. To decode neuronal spiking activity from the data, algorithms are first required to detect the location of cells in the video. It is still common for scientists to perform this task manually, as the heterogeneity in cell shape and frequency of cellular overlap impede automatic segmentation algorithms. We developed a versatile algorithm based on a popular image segmentation approach (the Level Set Method) and demonstrated its capability to overcome these challenges. We include no assumptions on cell shape or stereotypical temporal activity. This lends our framework the flexibility to be applied to new datasets with minimal adjustment.

## 1. Introduction

Two-photon calcium imaging has enabled the long-term study of neuronal population activity during learning and behaviour (Peron et al., 2015b). State of the art genetically encoded calcium indicators have sufficient signal-to-noise ratio (SNR) to resolve single action potentials (Chen et al., 2013). Furthermore, recent developments in microscope design have extended the possible field-of-view in which individual neurons can be resolved to 9.5mm^2^ (Stirman et al., 2016), and enabled the simultaneous imaging of separate brain areas (Lecoq et al., 2014). However, a comprehensive study of activity in even one brain area can produce terabytes of imaging data (Peron et al., 2015a), which presents a considerable signal processing problem.

To decode spiking activity from imaging data, one must first be able to accurately detect regions of interest (ROIs), which may be cell bodies, neurites or combinations of the two. Heterogeneity in the appearance of ROIs in imaging datasets complicates the detection problem. The calcium indicator used to generate the imaging video affects both a cell’s resting fluorescence and its apparent shape. For example, some genetically encoded indicators are excluded from the nucleus and therefore produce fluorescent ‘donuts’. Moreover, imaging data is frequently contaminated with measurement noise and movement artefacts. These challenges necessitate flexible, robust detection algorithms with minimal assumptions on the properties of ROIs.

Manual segmentation of calcium imaging datasets is still commonplace. While this allows the use of complex selection criteria, it is neither reproducible nor scalable. To incorporate implicitly a human’s selection criteria, which can be hard to define mathematically, supervised learning from extensive human-annotated data has been implemented (Valmianski et al., 2010; Apthorpe et al., 2016). Other approaches rely on more general cellular properties, such as their expected size and shape (Ohki et al., 2005) and that they represent regions of peak local correlation (Smith and Häusser, 2010; Kaifosh et al., 2014). The latter approaches use lower-dimensional summary statistics of the data, which reduces computational complexity but does not typically allow detection of overlapping regions.

To better discriminate between neighbouring cells, some methods make use of the temporal activity profile of imaging data. The (2+1)-D imaging video, which consists of two spatial dimensions and one temporal dimension, is often prohibitively large to work on directly. One family of approaches therefore reshapes the (2+1)-D imaging video into a 2-D matrix. The resulting matrix admits a decomposition — derived from a generative model of the imaging video — into two matrices, each encoding spatial and temporal information. The spatial and temporal components are estimated using a variety of methods, such as independent component analysis (Schultz et al., 2009; Mukamel et al., 2009) or non-negative matrix factorization (Maruyama et al., 2014). Recent variants extend the video model to incorporate detail on the structure of neuronal intracellular calcium dynamics (Pnevmatikakis et al., 2016) or the neuropil contamination (Pachitariu et al., 2016). By expressing the (2+1)-D imaging video as a 2-D matrix, this type of approach can achieve high processing speeds. This does, however, come at the cost of discarded spatial information, which can necessitate post-processing with morphological filters (Pnevmatikakis et al., 2016; Pachitariu et al., 2016).

In this paper, we propose a method in which cell boundaries are detected by multiple coupled active contours. To evolve an active contour we use the level set method, which is a popular tool in bioimaging due to its topological flexibility (Delgado-Gonzalo et al., 2015). To each active contour, we associate a higher-dimensional function, referred to as the level set function, whose zero level set encodes the contour location. We implicitly evolve an active contour via the level set function. The evolution of the level set function is driven by a local model of the imaging data temporal activity. The data model includes no assumptions on a cell’s morphology or stereotypical temporal activity. Our algorithm is therefore versatile, it can be applied to a variety of data types with minimal adjustment. For convenience, we refer to our method as ABLE (an Activity-Based LEvel set method). In the following, we describe the method and demonstrate its versatility and robustness on a range of *in vitro* and *in vivo* datasets.

## 2. Materials & Methods

### 2.1 Estimating the boundary of an isolated cell

Consider a small region of a video containing one cell (e.g. inside the dashed box, Fig. 1A). This region is composed of two subregions: the cell and the background. We want to partition the region into Ω^in^ and Ω^ou^^t^, where Ω^in^ corresponds to the cell and Ω^out^ the background. We compute a feature of the respective subregions, f ^in^ and f ^out^, with which to classify pixels into the cell interior or background. In particular, we define f ^in^ *∈* ℝ^*T*^ and f ^out^ *∈* ℝ^*T*^ as the average subregion time courses, where *T* is the number of frames in the video. We estimate the optimal partition as the one that minimizes discrepancies between a pixel’s time course and the average time course of the subregion to which it belongs. To calculate this discrepancy, we employ a dissimilarity metric, *D* (see below), which is identically zero when the time courses are perfectly matched and positive otherwise. As such, we minimize the following cost function, which we refer to as the external energy,

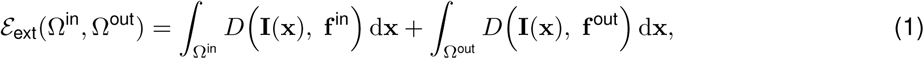

where *I*(x) *∈* ℝ^*T*^ is the time course of pixel x.

**Figure 1:**
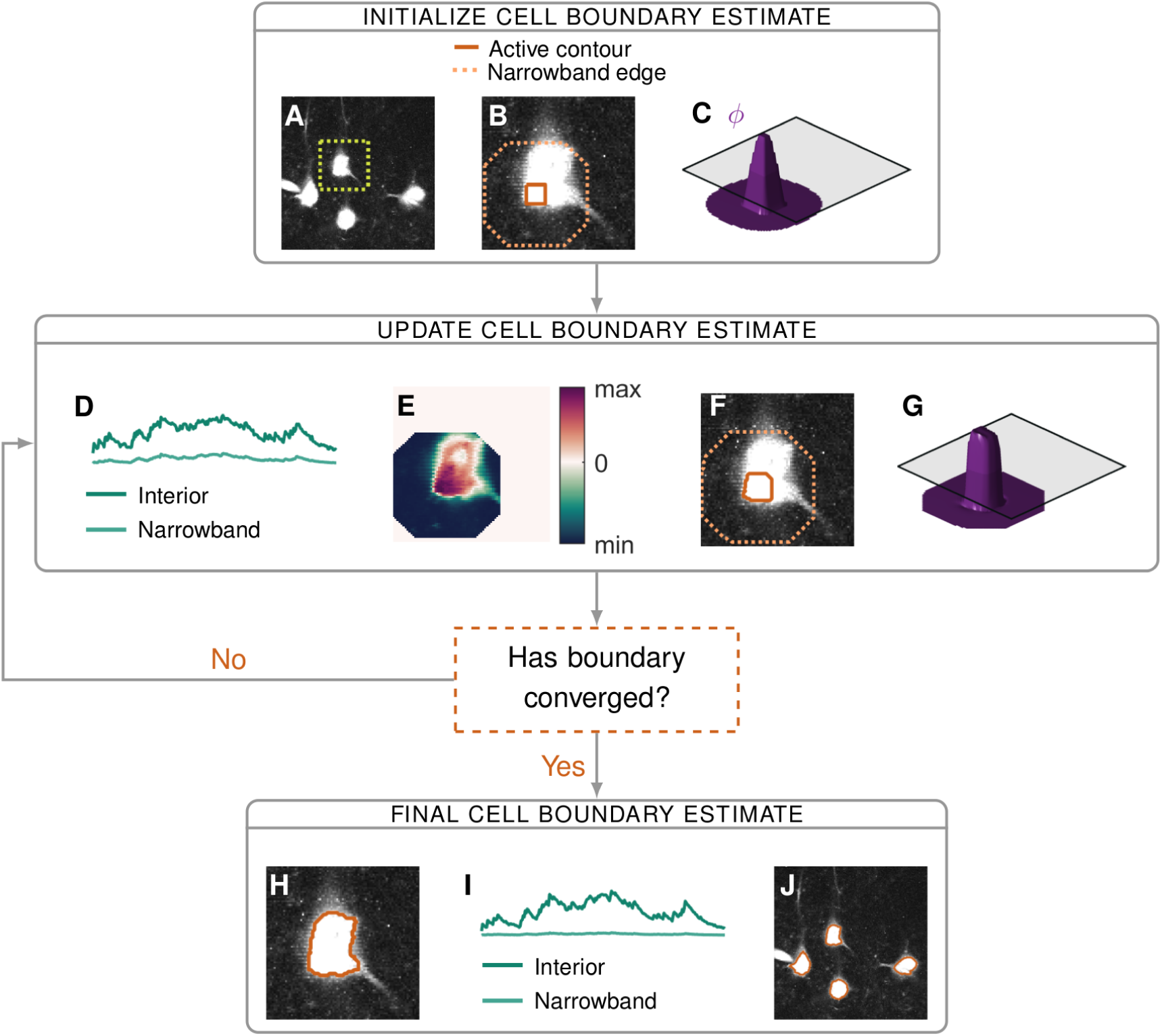
A flow diagram of the main steps of the proposed segmentation algorithm: initialization (**A**-**C**), iterative updates of the estimate (**D**-**G**) and convergence (**H**-**J**). When cells are sufficiently far apart we can segment them independently — in this example we focus on the isolated cell in the dashed box on the maximum intensity image in **A**. We make an initial estimate of the cell interior, from which we form the corresponding narrowband (**B**) and level set function *φ* (**C**). Based on the discrepancy between a pixel’s time course and the time courses of the interior and narrowband regions (**D**), we calculate the velocity of *φ* at each pixel (**E**). *φ* evolves according to this velocity (**G**), which updates the location of the interior and narrowband (**F**). Final results for: one cell (**H**), the average signals from the corresponding interior and narrowband (**I**) and segmentation of all four cells (**J**).

The cell location estimate is iteratively updated by the algorithm. At each iteration, the cell exterior is defined as the set of pixels within a fixed distance of the current estimate of the cell interior, see Fig. 1B. The default distance is taken to be two times the expected radius of a cell. We refer to this exterior region as the narrowband to emphasise its proximity to the contour of interest. The boundary between the interior and the narrowband is the active contour. As an active contour is updated, so is the corresponding narrowband (Fig. 1F). The region of the video for which the optimal partition is sought is therefore not static; rather, it evolves as an active contour evolves.

### 2.2 Computing the dissimilarity metric

Due to the heterogeneity of calcium imaging data, we do not use a universal dissimilarity metric. When both the pattern and magnitude of a pixel’s temporal activity are informative, as is typically the case for synthetic dyes, we use a measure based on the Euclidean distance, where

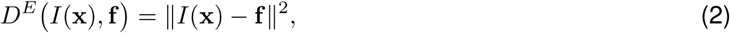

for f *∈* ℝ^*T*^. When we have an image not a video (i.e. **I**(x) and **f** are one-dimensional) this dissimilarity metric reduces to the fitting term introduced by Chan and Vese (2001). For datasets in which the fluorescence expression level varies significantly throughout cells and, as a consequence, pixels in the same cell exhibit the same pattern of activity at different magnitudes, we use a measure based on the correlation, such that

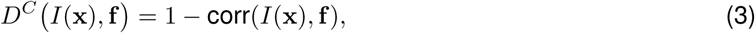

where corr represents the Pearson correlation coefficient. In this paper, as default, we use the Euclidean dissimilarity metric. Additionally, we present two notable examples in which the correlation-based metric is preferable.

### 2.3 External energy for neighbouring cells

We now extend the cost function presented in Eq. (1) to one suitable for partitioning a region into multiple cell interiors, {Ω^in,1^, Ω^in,2^*, …,* Ω^in*,M*^}, and a global exterior, Ω^out^, which encompasses the narrowbands of all the cells. We denote with f ^in*,i*^ the average time course of pixels exclusively in Ω^in*,i*^. Due to the relatively low axial resolution of a two-photon microscope, fluorescence intensity at one pixel can originate from multiple cells in neighbouring z-planes. Accordingly, we allow cell interiors to overlap when this best fits the data. In particular, we assume that a pixel in multiple cells would have a time course well fit by the sum of the interior time courses for each cell. The external energy in the case of multiple cells is thus

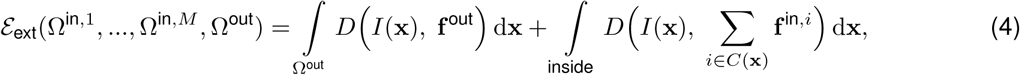

where the area termed ‘inside’ denotes the union of all cell interiors and the function *C*(**x**) identifies all cells whose interior contains pixel **x**. When the region to be partitioned contains only one cell, the external energy in Eq. (4) reduces to that in Eq. (1).

### 2.4 Level Set Method

It is not possible to find an optimal cell boundary by minimizing the external energy directly (Chan and Vese, 2001). An alternative solution is to start from an initial estimate, see below, and evolve this estimate in terms of an evolution parameter. *τ* In this approach, the boundary is called an active contour. To update the active contour we use the Level Set Method of Osher and Sethian (1988). This method was first introduced to image processing by Caselles et al. (1993) and Malladi et al. (1995); it has since found widespread use in the field. We implicitly represent the evolving boundary estimate of the *i*^th^ cell — the *i*^th^ active contour — by a function *φ*_*i*_, where *φ*_*i*_ is positive for all pixels in the cell interior, negative for those in the narrowband and zero for all pixels on the boundary (see Fig. 1C). We refer to *φ*_*i*_ as a level set function, as its zero level set identifies the contour of interest. We note that since the contour evolves with *τ*, *φ*_*i*_ itself depends upon *τ*. In the following, we present a set of *M* partial differential equations (PDEs) — one for each active contour — derived in part from Eq. (4), which dictate the evolution of the level set functions. The solution to the set of PDEs yields (as the zero level sets) the cell boundaries which minimize the external energy in Eq. (4).

From the external energy and a regularization term (Li et al., 2010), we define a new cost function

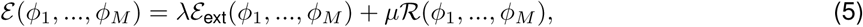

where the arguments to the external energy in Eq. (4) are replaced by the corresponding level set functions. The parameters *λ* and *μ* are real-valued scalars, which define the relative weight of the external energy and the regularizer. The regularizer is designed to ensure that a level set function varies smoothly in the vicinity of its active contour. The corresponding regularization energy is minimised when *φ*_*i*_ has gradient of magnitude one near the active contour and magnitude zero far from the contour. An example of such a function, a signed distance function (which is the shape of all level set functions upon initialization), can be seen in Fig. 1C.

A standard way to obtain the level set function that minimizes the cost function is to find the steady-state solution to the gradient flow equation (Aubert and Kornprobst, 2006), we do this for each *φ*_*i*_:

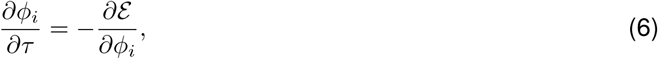

for *i ∈* {1, 2*, …, M*}. From Eq. (5) we obtain

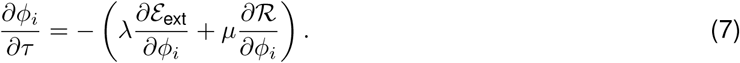

We solve this PDE numerically, by discretizing the evolution parameter *τ*, such that

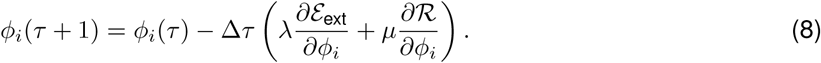

The regularization term, which encourages *φ*_*i*_ to vary smoothly in the image plane, helps to ensure the accurate computation of the numerical solution.

At every timestep *τ*, each level set function is consecutively updated until convergence. We must retain *μ*Δ*τ <* 0.25 in order to satisfy the Courant-Friedrichs-Lewy condition (Li et al., 2010) — a necessary condition for the convergence of a numerically-solved PDE. This condition requires that the numerical waves propagate at least as fast as the physical waves (Osher and Fedkiw, 2003). We therefore set Δ*τ* = 10 and *μ* = 0.2*/*Δ*τ*. For each dataset, we tune the value of *λ* based on the algorithm performance on a small section of the video. To attain segmentation results on the real datasets presented in this paper, we use *λ* = 150 (Section 3.1), *λ* = 50 (Section 3.3), *λ* = 25 (Section 3.4) and *λ* = 10 (Section 3.5).

### 2.5 External velocity

The movement of a level set function, *φ*_*i*_, is driven by the derivatives in Eq. (8)— *∂E*_ext_*/∂φ*_*i*_ provides the impetus from the video data and *∂R/∂φ*_*i*_ the impetus from the regulariser. In the following, we outline the calculation and interpretation of *∂E*_ext_*/∂φ*_*i*_; the regulariser is standard and its derivative is detailed in Li et al. (2010). As is typical in the level set literature (Zhao et al., 1996; Li et al., 2010), using an approximation of the Dirac delta function *δ*_*∊*_, we obtain an approximation of the derivative: *∂E*_ext_*/∂φ*_*i*_(x) = *δ*_*∊*_ (*φ*_*i*_(x)) *V*_*i*_(x), where

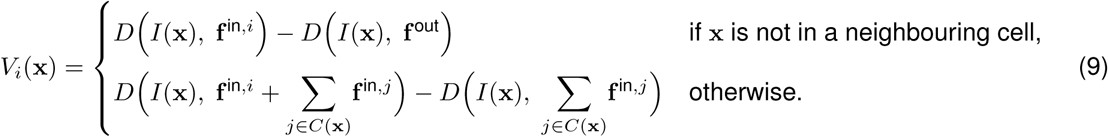

We refer to *V*_*i*_(**x**) as the external velocity as it encapsulates the impetus to movement derived from the external energy in Eq. (4), see Fig. 1E for an illustrative example.

The term *δ*_*∊*_, which is only non-zero at pixels on or near the cell boundary, acts as a localization operator, ensuring that the velocity only impacts *φ*_*i*_ at pixels in the vicinity of the active contour. The parameter *∊* defines the approximate radius, in pixels, of the non-zero band — here, we take *∊* = 2. The product with the localization operator means that, in practise, the external velocity must only be evaluated at pixels on or near the cell boundary. As a consequence, although the external velocity contains contributions from all cells in the video, the problem remains local — only neighbouring cells directly affect a cell’s evolution.

Although Ω^out^ represents a global exterior, in practise, we calculate the corresponding time course in Eq. (9), **f** ^out^, locally. To evaluate the external velocity of an active contour, we calculate **f** ^out^ as the average time course from pixels in the corresponding narrowband. This allows us to neglect components such as intensity inhomogeneity and neuropil contamination, see Fig. 2, which we assume vary on a scale larger than that of the narrowband.

**Figure 2:**
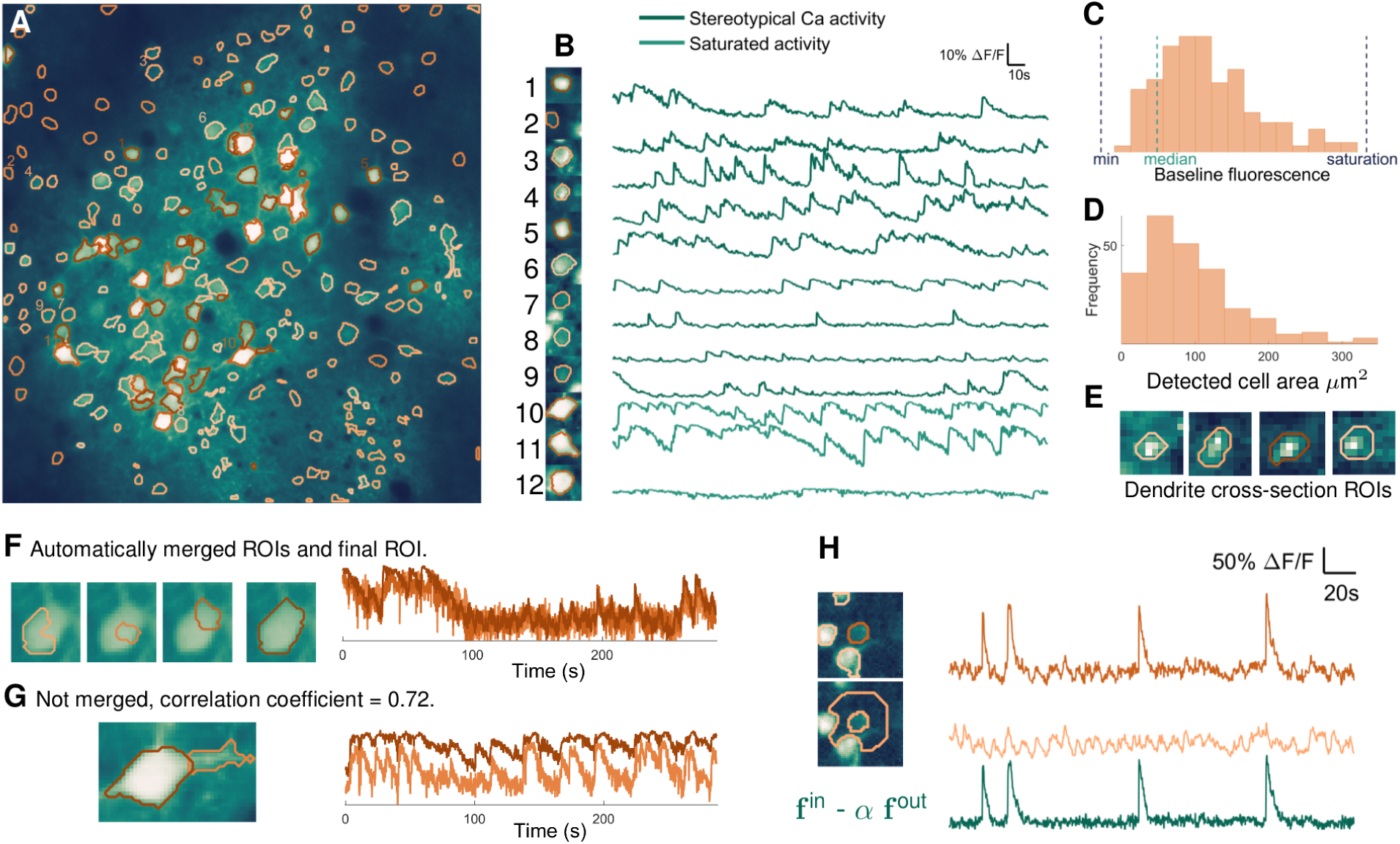
ABLE detects cells with varying size, shape and baseline intensity from mouse *in vivo* imaging data. The 236 detected ROIs are superimposed on the mean image of the imaging video (**A**). Extracted neuropil-corrected time series and corresponding ROIs are displayed for a subset of the detected regions (**B**). Cells with both stereotypical calcium transient activity (**B**, 1-9) and saturating fluorescence (**B**, 10-12) are detected. The performance of ABLE does not deteriorate due to intensity inhomogeneity: ROIs with baseline fluorescence from beneath the video median to just below saturation are detected (**C**). The area of detected regions varies (**D**) with the smallest ROIs corresponding to cross-sections of dendrites (**E**). Neighbouring regions with sufficiently high correlation are merged (**F**), those with lower correlation are not merged (**G**). In **F** we plot the ROIs prior to and after merging along with the corresponding neuropil-corrected time courses. In **G** we plot the separate ROIs and the neuropil-corrected time courses. The proposed method naturally facilitates neuropil-correction — the removal of the weighted, local neuropil time course from the raw cellular time course (**H**).

The external velocity of a single active contour, Eq. (9), can be interpreted as follows: if a pixel, not in another cell, has time course more similar to that of the contour interior than the narrowband, then the contour moves to incorporate that pixel. If a pixel in another cell has time course better matched by the sum of the interior time courses of cells containing that pixel plus the interior time course of the evolving active contour, then the contour moves to incorporate it. Otherwise, the contour is repelled from that pixel.

### 2.6 Initialization

We devised an automatic initialization algorithm which selects connected areas of either peak local correlation or peak mean intensity as initial ROI estimates. Initializing areas of peak mean intensity, which may correspond to artefacts rather than active cells (see e.g. the electrode in Fig. 1A), is essential so that these regions do not distort the narrowband signal of another ROI. We first compute the correlation image of the video. For each pixel, this is the average correlation between that pixel’s time course and those of the pixels in its 8-connected neighbourhood. Local peaks in this image and the mean intensity image are identified (by a built-in MATLAB function, ‘imextendedmax’) as candidate ROIs. The selectivity of the initialization is set by a tuning parameter *α*, which defines the relative height with respect to neighbouring pixels (in units of standard deviation of the input image) of the peaks that are suppressed. The higher the value of *α*, the more conservative the initialization. We have found it best to use a low value for *α* (in the range 0.2 - 0.8) so as to overestimate the number of ROIs; redundant estimates are automatically pruned during the update phase of the algorithm. Moreover, smaller values of *α* produce smaller initializations, which reduce errors due to initializations composed of multiple cells.

On synthetic data with dimensions 512 × 512 *× T*, the runtime of ABLE (minutes) increases linearly with the number of cells and is not significantly affected by increasing number of frames, *T*. Runtime was measured on a PC with 3.4GHz Intel Core i7 CPU.

### 2.7 Convergence

We stop updating a contour estimate if a maximum number of iterations *N*_max_ has been reached or the active contour has converged — using one or both of these conditions is common in the active contour literature, see, for example, Delgado-Gonzalo and Unser (2013); Li et al. (2010). A contour is deemed to have converged if, in *N*_con_ consecutive iterations, the number of pixels that are added to or removed from the interior is less than *ρ*. As default, we take *N*_max_ = 100, *N*_con_ = 40 and *ρ* = 2.

The complexity of the level set method is intrinsically related to the dimensionality of the active contour; the number of frames of the video is only relevant to the evaluation of the external velocity, Eq. (9), which accounts for a small fraction of the computational cost. In Table 1, we demonstrate that increasing video length by a factor of 10 has only a minor impact on processing time. As the framework includes no assumptions on an ROI’s stereotypical temporal activity, prior to segmentation a video can be downsampled by averaging consecutive samples, thereby simultaneously enabling the processing of longer videos and increasing signal-to-noise ratio.

**Table 1:** Runtime (minutes) on synthetic data of size 512 × 512 × *T*.

|  |  | Num. cells |  |  |
| --- | --- | --- | --- | --- |
|  |  | 25 | 125 | 225 |
| Num. frames ($T$) | 100 | 1.1 | 6.5 | 11.2 |
|  | 1000 | 1.3 | 6.5 | 12.7 |

Increasing cell density principally impacts the calculation of the external velocity and does, therefore, not alter the computational complexity of the algorithm. On synthetic data, we observe that increasing cell density only marginally affects the convergence rate, see Table 2. As emphasised in Section 2.5, updating an active contour is a local problem — consequently, we observe that algorithm runtime increases linearly with the total number of cells, see Table 1. Due to the independence of spatially separate ROIs in our framework, further performance speed-ups are achievable by parallelizing the computation.

**Table 2:**
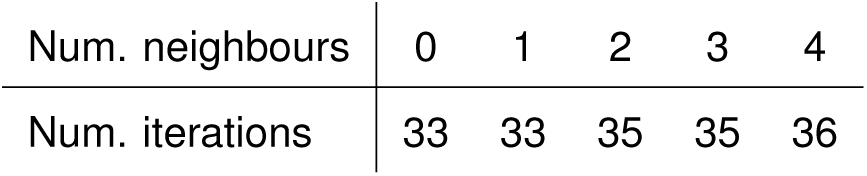
Number of iterations to convergence as cell density increases.

| Num. neighbours | 0 | 1 | 2 | 3 | 4 |
| --- | --- | --- | --- | --- | --- |
| Num. iterations | 33 | 33 | 35 | 35 | 36 |

On synthetic data the average number of iterations to convergence, over 100 realizations of noisy data, marginally increases as the number of cells in a given cell’s narrowband (‘neighbouring cells’) increases.

### 2.8 Merging and pruning ROIs

ABLE automatically merges two cells if they are sufficiently close and their interiors sufficiently correlated — a strategy previously employed in the constrained matrix factorization algorithm of Pnevmatikakis et al. (2016). When two contours are merged, their respective level set functions are replaced with a single level set function, initialized as a signed distance function (Fig. 1C), with a zero level set that represents the union of the contour interiors.

The required proximity for two cells to be merged is one cell radius (the expected cell radius is one of two required user input parameters). To determine the correlation threshold we consider the correlation of two noisy time courses corresponding to the average signals from two distinct sets of pixels belonging to the same cell. We assume the underlying signal components — which correspond to the cellular signal plus background contributions — have maximal correlation but that the additive noise reduces the correlation of the noisy time courses. Assuming the noise processes are independent from the underlying cellular signal and each other, the correlation coefficient of the noisy time courses is

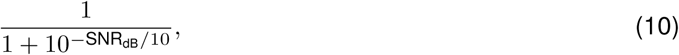

where SNR_dB_ is the signal-to-noise-ratio (dB) of the noisy time courses. We thus merge components with correlation above this threshold. We select a default correlation threshold of 0.8, derived from a default expected SNR of 5 (dB). The user has the option to input an empirically measured SNR, which updates the correlation threshold using the formula in Eq. (10).

A contour is automatically removed (‘pruned’) during the update phase if its area is smaller or greater than adjustable minimum or maximum size thresholds, which, as default, are set at 3 and 3*πr*^2^ pixels, respectively, where *r* is the expected radius of a cell.

### 2.9 Metric definitions

The signal-to-noise ratio (SNR) is defined as the ratio of the power of a signal ***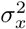*** and the power of the noise ***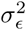***, such that ***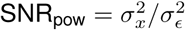***. We write the SNR in decibels (dB) as SNR_dB_ = 10 log10 (SNR_pow_).

Given two sets of objects, a ground truth set and a set of estimates, the precision is the percentage of estimates that are also in the ground truth set and the recall is the percentage of ground truth objects that are found in the set of estimates. As a complement of the precision we use the fall-out rate, the percentage of estimates not found in the ground truth set. The success rate (%) is

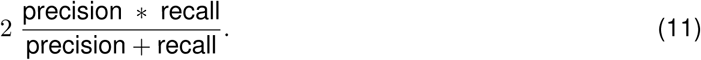

When the objects are cells, an estimate is deemed to match a ground truth cell if their centres are within 5 pixels of one another. When the objects are spikes, the required distance is 0.22s (3 sample widths). To quantify spike detection performance we also use the Root-Mean-Square-Error (RMSE), which is the square root of the average squared error between an estimated spike time 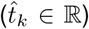 and the ground truth spike time (*t*_*k*_ *∈* ℝ).

### 2.10 Simulations

To quantify segmentation performance, we simulated calcium imaging videos. In the following, we detail the method used to generate the videos. Cellular spike trains are generated from mutually independent Poisson processes. A cell’s temporal activity is the sum of a stationary baseline component, the value of which is selected from a uniform distribution, and a spike train convolved with a stereotypical calcium transient pulse shape. Cells are ‘donut’ (annulus) shaped to mimic videos generated by genetically encoded calcium indicators, which are excluded from the nucleus. To achieve this, the temporal activity of a pixel in a cell is generated by multiplying the cellular temporal activity vector by a factor in [0, 1] that decreases as pixels are further from the cell boundary. When two cells overlap in one pixel, we sum the contributions of both cells at that pixel. Spatially and temporally varying background activity, generated independently from the cellular spiking activity, is present in pixels that do not belong to a cell.

### 2.10 Software accessibility

The software described in the paper is freely available online at http://github.com/StephanieRey.

### 2.12 Two-photon calcium imaging of quadruple whole-cell recordings

All procedures conformed to the standards and guidelines set in place by the Canadian Council on Animal Care, under the Research Institute of the McGill University Health Centre animal use protocol number 2011-6041 to PJS. P11-P15 mice of either sex were anaesthetised with isoflurane, decapitated, and the brain was rapidly dissected in 4°C external solution consisting of 125 mM NaCl, 2.5 mM KCl, 1 mM MgCl_2_, 1.25 mM NaH_2_PO_4_, 2 mM CaCl2, 26 mM NaHCO3, and 25 mM dextrose, bubbled with 95% O2/5% CO2 for oxygenation and pH. Quadruple whole-cell recordings in acute visual cortex slices were carried out at 32°C-34°C with internal solution consisting of 5 mM KCl, 115 mM K-gluconate, 10 mM K-HEPES, 4 mM MgATP, 0.3 mM NaGTP, 10 mM Na-phosphocreatine, and 0.1% w/v biocytin, adjusted with KOH to pH 7.2-7.4. On the day of the experiment, 20 *μ*M Alexa Fluor 594 and 180 *μ*M Fluo-5F pentapotassium salt (Life Technologies) were added to the internal solution. Electrophysiology amplifier (Dagan Corporation BVC-700A) signals were recorded with a National Instruments PCI-6229 board, using in-house software running in Igor Pro 6 (WaveMetrics). Two-photon excitation was achieved by raster-scanning a Spectraphysics MaiTai BB Ti:Sa laser tuned to 820 nm across the sample using an Olympus 40x objective and galvanometric mirrors (Cambridge Technologies 6215H, 3 mm, 1 ms/line, 256 lines). Substage photomultiplier tube signals (R3896, Hamatsu) were acquired with a National Instruments PCI-6110 board using ScanImage 3.7 running in MATLAB (MathWorks). Layer-5 pyramidal cells were identified by their prominent apical dendrites using infrared video Dodt contrast. Unless otherwise stated, all drugs were obtained from Sigma-Aldrich.

### 2.13 Two-photon calcium imaging of bulk loaded hippocampal slices

All procedures were performed in accordance with national and institutional guidelines and were approved by the UK Home Office under Project License 70/7355 to SRS. Juvenile wild-type mice of either sex (C57Bl6, P13-P21) were anaesthetised using isoflurane prior to decapitation procedure. Brain slices (400 *μ*m thick) were horizontally cut in 1-4°C ventilated (95% O2, 5% CO2) slicing Artificial Cerebro-Spinal Fluid (sACSF: 0.5 mM CaCl2, 3.0 mM KCl, 26 mM NaHCO3, 1 mM NaH2PO4, 3.5 mM MgSO4, 123 mM Sucrose, 10 mM D-Glucose). Hippocampal slices containing Dentate Gyrus, CA3 and CA1 were taken and resting in ventilated recovery ACSF (rACSF: 2 mM CaCl_2_, 123 mM NaCl, 3.0 mM KCl, 26 mM NaHCO_3_, 1mM NaH_2_PO_4_, 2mM MgSO_4_, 10mM D-Glucose) for 30min at 37°C. After this the slices were placed in an incubation chamber containing 2.5 mL of ventilated rACSF and ‘painted’ with 10 *μ*L of the following solution: 50 *μ*g of Cal-520 AM (AAT Bioquest), 2 *μ*L of Pluronic-F127 20% in DMSO (Life Technologies) and 48 *μ*L of DMSO (Sigma Aldrich) where they were left for 30 min at 37°C in the dark. Slices were then washed in rACSF at room temperature for 30 min before imaging. Dentate Gyrus granular cells were identified using oblique illumination prior to being imaged using a standard commercial galvanometric scanner based two-photon microscope (Scientifica Ltd) coupled to a mode-locked Mai Tai HP Ti Sapphire (Spectra-Physics) laser system operating at 810 nm. Functional calcium images of granular cells were acquired with a 40X objective (Olympus) by raster scanning a 180 × 180 *μ*m^2^ square Field of View at 10 Hz. Electrical stimulation was accomplished with a tungsten bipolar concentric microelectrode (WPI) where the tip of the electrode was placed into the molecular layer of the Dentate Gyrus (20 pulses with a pulse-width of 400 *μ*s and a 60 *μ*A amplitude were delivered into the tissue with a pulse repetition rate of 10 Hz, repeated every 40 sec). Unless otherwise stated, all drugs were obtained from Sigma-Aldrich.

## 3. Results

### 3.1 ABLE is robust to heterogeneity in cell shape and baseline intensity

ABLE detected 236 ROIs with diverse properties from the publicly available mouse *in vivo* imaging dataset of Peron et al. (2015c), see Fig. 2. Automatic initialization on this dataset produced 253 ROIs with 17 automatically removed during the update phase of the algorithm after merging with another region.

To maintain a versatile framework we included no priors on cellular morphology in the cost function that drives the evolution of an active contour. This allowed ABLE to detect ROIs with varied shapes (Fig. 2A) and sizes (Fig. 2D). The smaller detected ROIs correspond to cross-sections of dendrites (Fig. 2E), whereas the majority correspond to cell bodies. The topological flexibility of the level set method allows cell bodies and neurites to be segmented as separate (Fig. 2G) or connected (Fig. 2A) objects, depending on the correlation between their time courses. ABLE automatically merges neighbouring regions that are sufficiently correlated (Fig. 2F). Cell bodies and dendrites that are initialised separately and exhibit distinct temporal activity, however, are not merged. For example, the cell body and neurite in Fig. 2G were not merged as the cell body’s saturating fluorescence time course was not sufficiently highly correlated with that of the neurite.

Evaluating the external velocity, which drives an active contour’s evolution, requires only data from pixels in close proximity to the contour (see Section 2.5). This region has radius of the same order as that of a cell body. Background intensity inhomogeneity, caused by uneven loading of synthetic dyes or uneven expression of virally inserted genetically encoded indicators, tend to occur on a scale larger than this. On this dataset we show that, as a result of this local approach, ABLE is robust to background intensity inhomogeneity. This is illustrated by the wide range of baseline intensities of the detected ROIs (Fig. 2C), some of which are even lower than the video median.

No prior information on stereotypical neuronal temporal activity is included in our framework. Cells detected by ABLE exhibit both stereotypical calcium transient activity (Fig. 2B:1-9) and non-stereotypical activity (Fig. 2B:10-12), perhaps corresponding to saturating fluorescence, higher firing cell types such as interneurons, or non-neuronal cells.

The scattering of photons when imaging at depth can result in leakage of neuropil signal into cellular signal. To obtain decontaminated cellular time courses it is thus important to perform neuropil correction in a subsequent stage, once cells have been located. This involves computation of the decontaminated cellular signal by subtracting the weighted local neuropil signal from the raw cellular signal. As illustrated in Fig. 2H, the proposed method naturally facilitates neuropil correction, as it computes the required components as a by-product of the segmentation process (see Section 2.5). The appropriate value of the weight parameter varies depending on the imaging set-up (Peron et al. 2015a; Chen et al. 2013; Kerlin et al. 2010). We therefore do not include neuropil-correction as a stage of the algorithm, preferring instead to allow users the flexibility to choose the appropriate parameter in post-processing.

### 3.2 ABLE demixes overlapping cells

When imaging through scattering tissue, a two-photon microscope can have relatively low axial resolution (on the order of ten microns) in comparison to its excellent lateral resolution. As a consequence, the photons collected at one pixel can in some cases originate from multiple cells in a range of z-planes. For this reason, cells can appear to overlap in an imaging video (for an example, see Fig. 3E). It is crucial that segmentation algorithms can delineate the true boundary of ‘overlapping’ cells, which we refer to as ‘demixing’, so that the functional activity of each cell can be correctly extracted and analysed. In a set of experiments on real and simulated data, we demonstrated that ABLE can demix overlapping cells.

**Figure 3:**
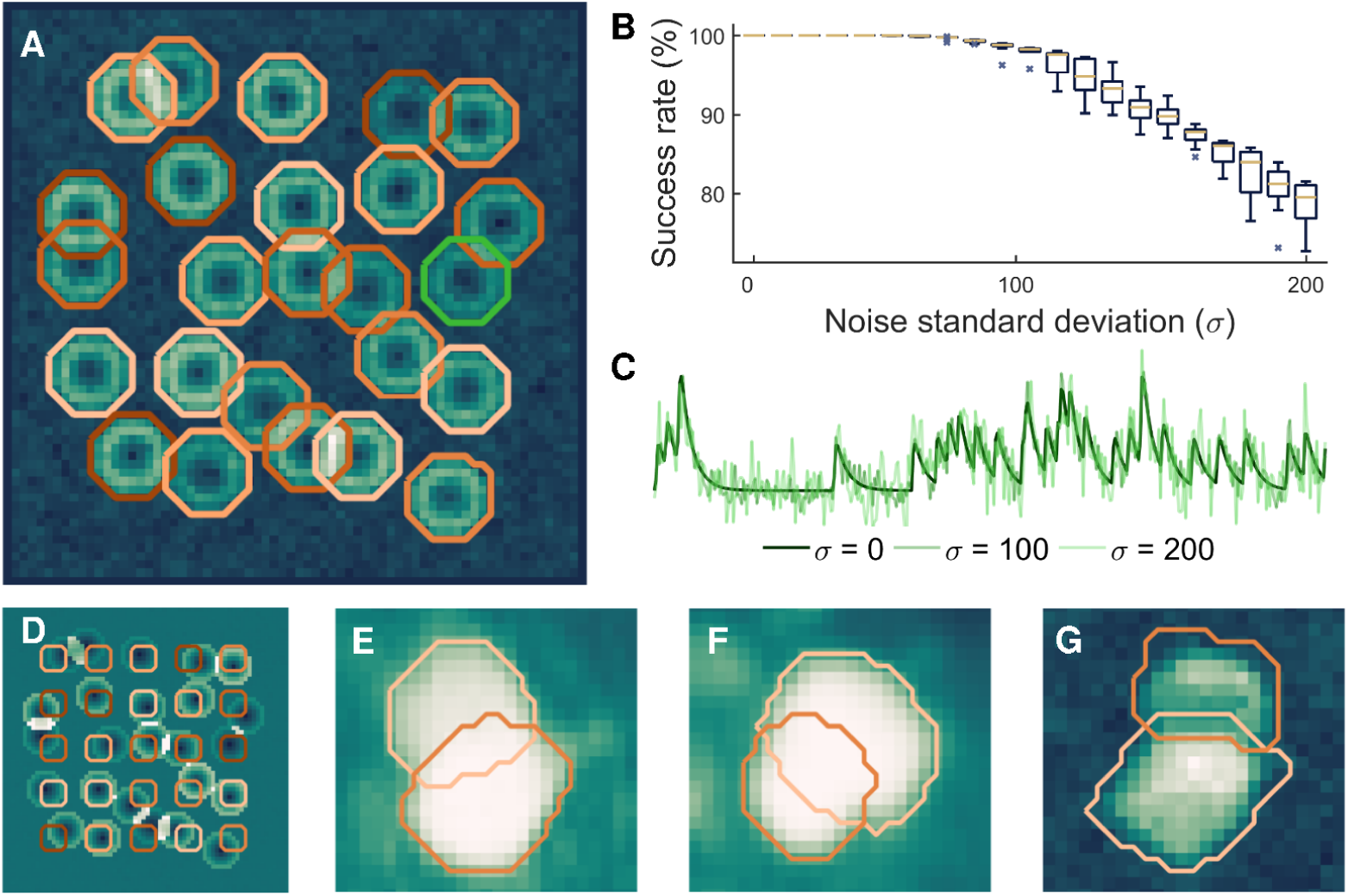
ABLE demixes overlapping cells in real and simulated data. With high accuracy, we detect the true boundaries of overlapping cells from noisy simulated data, the detected contours for one realization of noise with standard deviation (*σ*) 60 are plotted on the correlation image in **A**. Given an initialisation on a fixed grid, displayed on the mean image in **D**, we detect the true cell boundaries with success rate of at least 99% for *σ <* 90 (**B**). The central marker and box edges in **B** indicate the median and the 25^th^ and 75^th^ percentiles, respectively. For noise level reference, we plot the average time course from inside the green contour in **A** at various levels (**C**). ABLE demixes overlapping cells in real GCaMP6s mouse *in vivo* data — detected boundaries are superimposed on the mean image (**E** and **F**) and correlation image (**G**), respectively.

On synthetic data containing 25 cells, 17 of which had some overlap with another cell, we measured the success rate of ABLE’s segmentation compared to the ground truth cell locations (Fig. 3A-C), when the algorithm was initialised on a fixed grid (Fig. 3D). For full description of the performance metric used, see Section 2.9. Performance was measured over 10 realizations of noise at each noise level. On average, over all cells and noise realizations, ABLE achieved success rate greater than 99% when the noise standard deviation was less than 90 (Fig. 3B). Cells were simulated with uneven brightness to mimic the ‘donut’ cells generated by some genetically encoded indicators that are excluded from the nucleus. Consequently, the correlation-based dissimilarity metric was used on this data. As a result, pixels with significantly different resting fluorescence, but identical temporal activity pattern, were segmented in the same cell (Fig. 3A).

On the publicly available mouse *in vivo* imaging dataset of Peron et al. (2015c), ABLE demixed overlapping cells (Fig. 3E-F). In this dataset, the vibrissal cortex was imaged at various depths, from layer 1 to deep layer 3, whilst the mouse performed a pole localization task (Peron et al., 2015a; Guo et al., 2014). Some cells appear to overlap, due to the relatively low axial resolution when imaging at depth through tissue. When an ROI was initialised in each separate neuron, ABLE accurately detected the overlapping cell boundaries using the Euclidean distance dissimilarity metric, Eq. (2). On the Neurofinder Challenge dataset presented in Section 3.4, ABLE demixed overlapping cells when performing segmentation with the correlation-based dissimilarity metric, Eq. (3), see Fig. 3G.

### 3.3 ABLE detects synchronously spiking, densely packed cells

ABLE detected 207 ROIs from mouse *in vitro* imaging data (Fig. 4). Cells in this dataset exhibit activity that is highly correlated with other cells and the background as the brain slice was electrically stimulated (at rate 10Hz for 2s every 40s) during imaging. When the cell interior and narrowband time courses are highly correlated, the external velocity of the active contour, Eq. (9), derived from the Euclidean distance dissimilarity metric, Eq. (2), is driven by the discrepancy between the baseline intensities of the subregions. This is evident when we consider the average time course of the cell interior (f ^in^) and exterior (f ^out^) as a sum of a stationary baseline component — the resting fluorescence — and an activity component that is zero when a neuron is inactive, such that f ^in^ = *b*^in^ + a^in^ and f ^out^ = *b*^out^ + a^out^. The time course of a pixel x is *I*(x) = *b***^x^** + a**^x^**. Substituting these expressions into Eq. (9), for pixels not in another cell, we obtain the external velocity***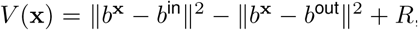***, where the residual, *R*, encompasses all terms with contributions from the activity components. When the cell and the background are highly correlated — meaning that the discrepancy between activity components is low and, consequently, the contribution from *R* is comparatively small — the external velocity will drive the contour to include pixels with baselines more similar to the interior than the background. As a result of this, ABLE detected ROIs despite their high correlation with the background (Fig. 4C). Furthermore, inactive ROIs were detected (Fig. 4 H-J), when their baseline fluorescence allowed them to be identified from the background (Fig. 4I).

**Figure 4:**
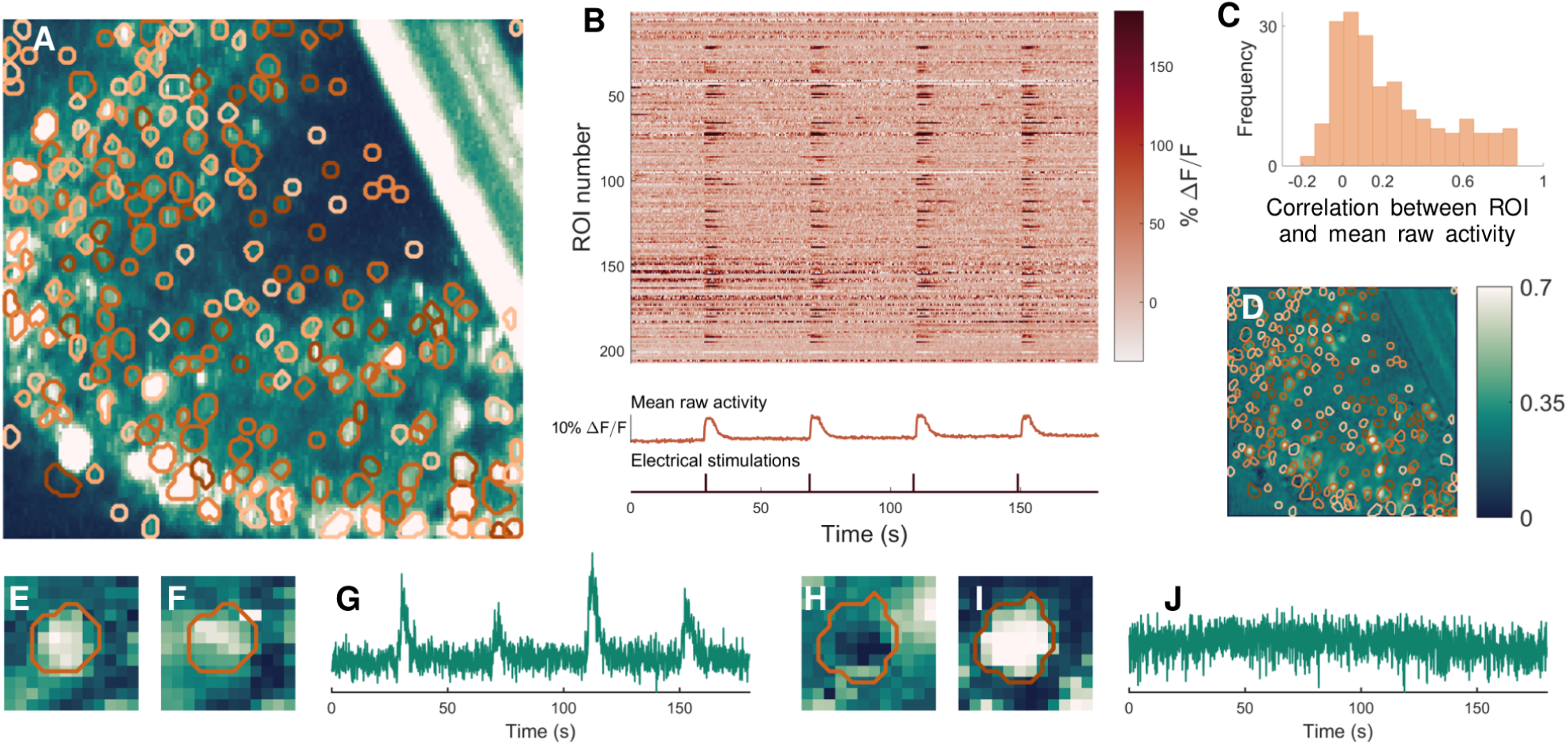
ABLE detects synchronously spiking, densely packed cells from mouse *in vitro* imaging data. The boundaries of the 207 detected ROIs are superimposed on the thresholded maximum intensity image (**A**) and the correlation image (**D**). For all correlation data we use Pearson’s correlation coefficient. ABLE detects ROIs that exhibit high correlation with the background **C** and neighbouring synchronously spiking ROIs (**B**). Panel **B** displays the neuropil-corrected extracted time courses of the 207 ROIs (each plotted as a row of the matrix) along with the video mean raw activity and the time points of the electrical stimulations. Panel **C** displays the histogram of the correlation coefficient between the mean raw activity of the video and the extracted time series of each ROI. ABLE detected both active (**E**-**G**) and inactive ROIs (**H**-**I**). We display the contours of the two detected ROIs on the correlation image (**E** and **H**), the mean image (**F** and **I**) and the corresponding extracted time courses (**G** and **J**).

The algorithm was automatically initialised on this dataset with 250 ROIs, initializations in the bar (an artefact that can be seen in the top right of Fig. 4A) were prohibited. Of the initialised ROIs, 19 were pruned automatically during the update phase of the algorithm as (i) their interior time course was not sufficiently different from that of the narrowband (3 ROIs), (ii) they merged with another region (2 ROIs) or (iii) they crossed the minimum and maximum size thresholds (14 ROIs).

### 3.4 Algorithm comparison on manually labelled dataset

We compared the performance of ABLE with two state of the art calcium imaging segmentation algorithms — CNMF (Pnevmatikakis et al., 2016) and Suite2p (Pachitariu et al., 2016) — on a manually labelled dataset from the Neurofinder Challenge, see Fig. 5. The dataset, which can be accessed at the Neurofinder Challenge website (see references), was recorded at 8Hz and generated using the genetically encoded calcium indicator GCaMP6s. Consequently, we apply ABLE with the correlation-based dissimilarity metric, Eq. (3), which is well suited to neurons with low baseline fluorescence and uneven brightness. As the dataset is large enough (512x512x8000 pixels) to present memory issues on a standard laptop, we run the patch-based implementation of CNMF, which processes spatially-overlapping patches of the dataset in parallel. We optimise the performance of each algorithm by selecting a range of values for each of a set of tuning parameters and generating segmentation results for all combinations of the parameter set. The results are visualised on the correlation image and the parameter set that presents the best match to the correlation image is selected. This process is representative of what a user may do in practise when applying an algorithm to a new dataset.

**Figure 5:**
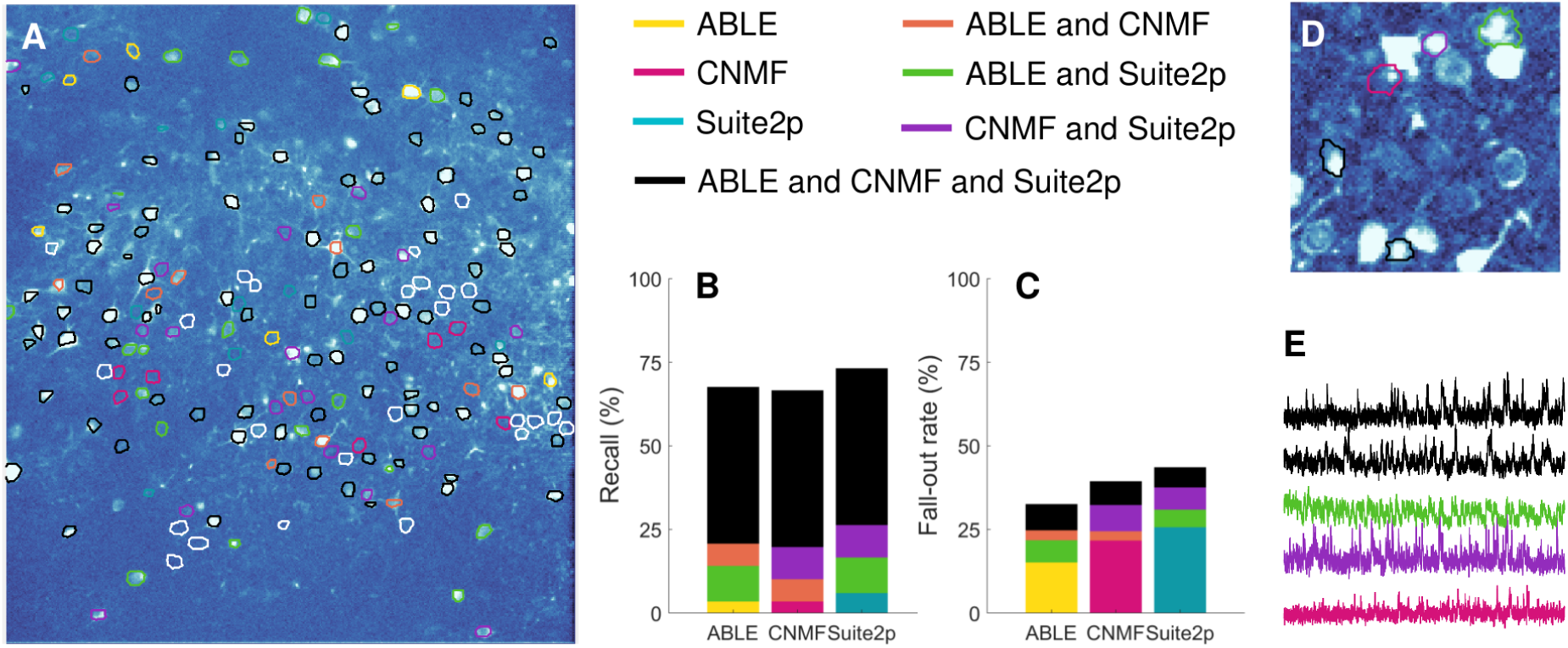
We compare the segmentation results of ABLE, CNMF (Pnevmatikakis et al., 2016) and Suite2p (Pachitariu et al., 2016) on a manually labelled dataset from the Neurofinder Challenge. On the correlation image we plot the boundaries of the manually labelled cells colour-coded by the combination of algorithms that detected them (**A**), undetected cells are indicated by a white contour. Suite2p detected the highest proportion of manually labelled cells (**B**), whereas ABLE had the lowest fall-out rate (**C**), which is the percentage of detected regions not present in the manual labels. Some algorithm-detected ROIs that were not present in the manual labels are detected by multiple algorithms (**D**) and have time courses which exhibit stereotypical calcium transient activity (**E**). The correlation image in **D** is thresholded to enhance visibility of local peaks in correlation. In **E**, we plot the extracted time courses of the ROIs in **D**.

ABLE achieved the highest success rate (67.5%) when compared to the manual labels, see Table 3. For a definition of the success rate and other performance metrics used, see Section 2.9. ABLE achieved a lower fall-out rate than Suite2p and CNMF (Fig. 5C) — 67.5% of the ROIs it detected matched with the manually labelled cells. Some of the ‘false positives’ were consistent among algorithms (Fig. 5C) and corresponded to local peaks in the correlation image (Fig. 5D), whose extracted time courses displayed stereotypical calcium transient activity (Fig. 5E). A subset of these ROIs may thus correspond to cells omitted by the manual operator. The highest proportion of the manually labelled cells were detected by Suite2p, which detected the greatest number of cells not detected by any other algorithm (Fig. 5B). A small proportion (13.2%) of cells were detected by none of the algorithms. As can be seen from Fig. 5A, these do not correspond to peaks in the correlation image, and may reflect inactive cells detected by the manual operator.

**Table 3:**
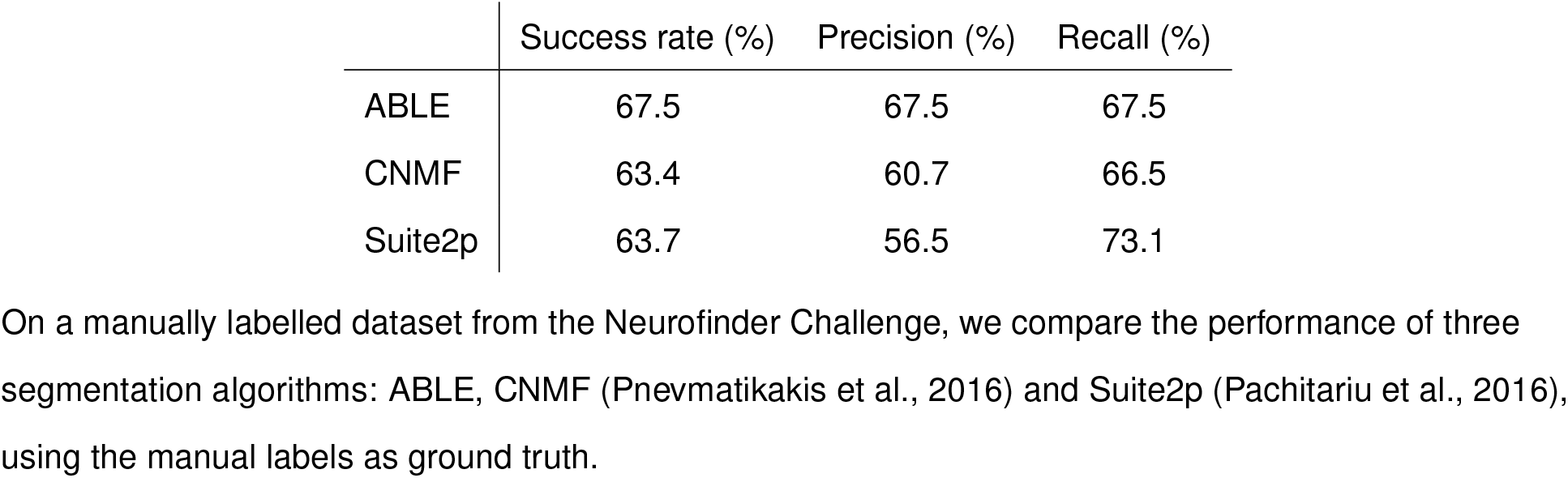
Algorithm success rate (1dp) on manually labelled dataset.

### 3.5 Spikes are detected from ABLE-extracted time courses with high temporal precision

Typically, after cells have been identified in calcium imaging data, spiking activity is detected from the extracted cellular time courses and the relationship between cellular activity (and, if measured, external stimuli) is analysed. On a mouse *in vitro* dataset (21 videos, each 30s long), we demonstrated that time courses from cells automatically segmented by ABLE allow spikes to be detected accurately and with high temporal precision (Fig. 6). The dataset has simultaneous electrophysiological recordings from four cells (the electrodes can be seen in the mean image Fig. 6A), which enabled us to compare inferred spike times from the imaging data with the ground truth. We performed spike detection automatically with an existing algorithm (Oñativia et al., 2013; Reynolds et al., 2016). On average, over all cells and recordings, 78% of ground truth spikes are detected with a precision of 88% (Fig. 6D). The error in the location of detected spikes is less than one sample width — the average absolute error was 0.053 (s).

**Figure 6:**
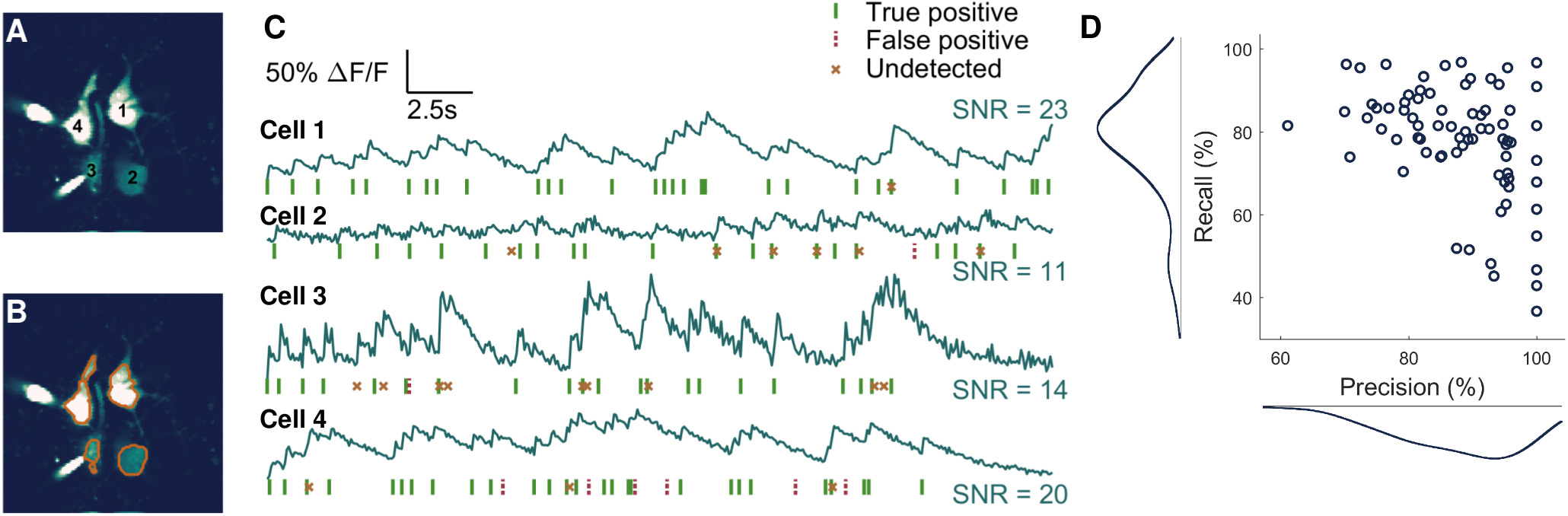
Spikes are detected from ABLE-extracted time courses with high accuracy. On an *in vitro* dataset (21 imaging videos, each 30s long) we demonstrate spike detection performance compared to electrophysiological ground truth on time courses extracted from cells segmented by ABLE. We plot the labelled cells (**A**) and corresponding boundaries detected by ABLE (**B**) on the mean image of one imaging video. The extracted cellular time courses and detected spikes are plotted in **C**. Spike detection was performed with an existing algorithm (Oñativia et al., 2013; Reynolds et al., 2016). On average over all videos, 78% of spikes are detected with a precision of 88% **D**.

## 4. Discussion

In this paper, we present a novel approach to the problem of detecting cells from calcium imaging data. Our approach uses multiple coupled active contours to identify cell boundaries. The core assumption is that the local region around a single cell (e.g. inside the dashed box Fig. 1A) can be well-approximated by two subregions, the cell interior and exterior. The average time course of the respective subregions is used as a feature with which to classify pixels into either subregion. We assume that pixels in which multiple cells overlap have time courses that are well-approximated by the sum of each cell’s time course. We form a cost function based on these assumptions that is minimised when the active contours are located at the true cell boundaries. Our results on real and simulated data indicate that this is a versatile and robust framework for segmenting calcium imaging data.

The cost function in our framework (Eq. 4) penalises discrepancies between the time course of a pixel and the average time course of the subregion to which it belongs. To calculate this discrepancy we use one of two dissimilarity metrics: one based on the correlation, which compares only patterns of temporal activity, the other based on the Euclidean distance, which implicitly takes into account both pattern and magnitude of temporal activity. When the latter metric is used, our cost function is closely related to that of Chan and Vese (2001). If we were to take as an input one frame of a video (or a 2D summary statistic such as the mean image), the external energy in our cost function for an isolated cell would be identical to the fitting term of Chan and Vese (2001). The lower-dimensional approach is, however, not sufficient for segmenting cells with neighbours that have similar baseline intensities. By incorporating temporal activity we can accurately delineate the boundaries of neighbouring cells (Fig. 3A).

We evolve one active contour for each cell identified in the initialization. Contours are evolved predominantly independently, with the exception of those within a few pixels of another active contour (see Section 2.5). In contrast to previous approaches to coupling active contours (Zimmer and Olivo-Marin, 2005; Dufour et al., 2005), we do not penalise overlap of contour interiors. This is because low axial resolution when imaging through scattering tissue can result in the signals of multiple cells being expressed in one pixel. We therefore permit interiors to overlap when the data is best fit by the sum of average interior time courses. Using this method we can accurately demix the contribution of multiple cells from single pixels in real and simulated data (Fig. 3).

ABLE is a flexible method: we include no priors on a region’s morphology or stereotypical temporal activity. Due to this versatility, ABLE segmented cells with varying size, shape, baseline intensity and cell type from a mouse *in vivo* dataset (Fig. 2). Moreover, only 2 parameters need to be set by a user for a new dataset. These are the expected radius of a cell and *λ*, the relative strength of the external velocity compared to the regulariser, see Eq. (5). In order to permit ABLE to segment irregular shapes such as cell bodies attached to dendritic branches (Fig. 2A), the weighting parameter, *λ*, must be set sufficiently high to counter the regulariser’s implicit bias towards smooth contours.

Unlike matrix factorization (Maruyama et al., 2014; Pnevmatikakis et al., 2016) and dictionary learning (Diego Andilla and Hamprecht, 2014), which fit a global model to an imaging video, our approach requires only local information to evolve a contour. To evolve an active contour, ABLE uses temporal activity from an area around that contour with size on the order of the radius of a cell. This allows us to omit from our model the spatial variation of the neuropil signal and baseline intensity inhomogeneities, which we assume to be constant on our scale. Our local approach means that the algorithm is readily parallelizable and, in the current implementation, runtime is virtually unaffected by video length (Table 1) and increases linearly with the number of cells.

Like any level set method, the performance of ABLE is bounded by the quality of the initialization — if no seed is placed in a neuron it will not be detected, if a seed is spread across multiple neurons they may be jointly segmented. In this work, we developed an automatic initialization algorithm that selects local peaks in the correlation and mean images as candidate ROIs. This approach, however, can lead to false negatives in dense clusters of cells in which the correlation image can appear smooth. In future work, an initialization based on temporal activity, rather than a 2D summary statistic, could overcome this issue. Our algorithm included minimal assumptions about the objects to be detected. To tailor ABLE to a specific data type (e.g. somas versus neurites), it is possible to incorporate terms relating to a region’s morphology or stereotypical temporal activity into the cost function. Furthermore, the level set method is straightforward to extend to higher dimensions (Dufour et al., 2005), which means our framework could be adapted to detect cells in light-sheet imaging data (Ahrens et al., 2013).

Here we have presented a framework in which multiple coupled active contours detect the boundaries of cells from calcium imaging data. We have demonstrated the versatility of our framework, which includes no priors on a cell’s morphology or stereotypical temporal activity, on real *in vivo* imaging data. In this data, we are able to detect cells of various shapes, sizes, and types. We couple the active contours in a way that permits overlap when this best fits the data. This allows us to demix overlapping cells on real and simulated data, even in high noise scenarios. Our results on a diverse array of real datasets indicate that ours is a flexible and robust framework for segmenting calcium imaging data.

